# Extracellular HDAC6 ZnF UBP domain enhances podosome-mediated neuronal migration

**DOI:** 10.1101/2024.12.04.626746

**Authors:** Tazeen Qureshi, Smita Eknath Desale, Shweta Kishor Sonawane, Abhishek Ankur Balmik, Hariharakrishnan Chidambaram, Subashchandrabose Chinnathambi

## Abstract

The increasing prevalence of Alzheimer’s disease demands research into therapeutic strategies that go beyond Amyloid and Tau. Synaptic loss, neuritic loss, and microtubule destabilization due to misfolded Tau and dysfunctional signalling in the AD-affected neuron, all point toward understanding and targeting cytoskeletal dysregulation in AD. Here we present, an extensive study on the novel role of the ZnF UBP (Zinc Finger Ubiquitin Binding Protein) domain of HDAC6 (Histone Deacetylase 6) in actin remodelling in neurons. We have demonstrated this function through immunofluorescence colocalization analysis in actin-rich podosome structures. We have found that this HDAC6 domain induces increased localization of the actin polymerization proteins, Arp2 and WASP and the adaptor protein TKS5 in the podosome structures. We have also extended our work to understand the potential of this domain in enhancing the podosome-mediated migration of neuronal cells. It was thus established that HDAC6 ZnF UBP induces increased association of cytoskeletal proteins within the podosomes, conferring enhanced migration potential to neurons and presenting an interesting strategy to improve neuronal health.

## 1. INTRODUCTION

Alzheimer’s disease (AD) is the most progressive neurodegenerative disease, characterized by the amyloid-beta plaques on the extracellular side of the neuron and the deposits of misfolded Tau protein within the neurons, making AD one of the most important Tauopathies. The major symptom experienced by AD patients is memory loss, caused due to the rapid degeneration of neurons and synaptic connections, thus emphasizing the involvement of the cytoskeleton in neurodegeneration. The current therapeutic strategies work on just symptom management but efficient mechanisms must be developed to counter the issues at the molecular level ^1^. The development of neuronal extensions and connections helps in navigation, efficient transfer and information processing as well as for the overall maintenance of neuronal homeostasis. Growth cones are specialized structures in neurons, responsible for the formation of such extensions. These neuritic extensions are composed of protrusions of filopodia (filamentous actin) alternating with fan-shaped lamellipodia (branched actin). These structures harbour dynamic coordination of the various cytoskeletal proteins such as actin, tubulin and their associated proteins in a process called ‘actin remodelling’ ^2,3^. Actin remodelling is a highly dynamic activity involving the crosstalk between several proteins resulting in nucleation, branching and assembly of actin filaments leading to the formation of complex actin networks and structures. Podosomes are actin-rich structures, most commonly found at the periphery of migrating and invading cells such as transformed cells, fibroblasts, macrophages, etc. Podosomes are abundant in matrix metalloproteases, which help degrade the extracellular matrix and enable migration of the cell ^4^. In the nervous system, podosomes are found in the microglia, which serve to keep the brain environment free from pathogenic organisms ^5^. Microglia have been reported to be stimulated and to migrate in the presence of amyloid plaques as well as extracellular Tau deposits ^6^. Recent studies have also established that microglia degrade extracellular Tau with the help of podosomes and other migratory structures of the growth cone ^7^.

Histone Deacetylases (HDACs) are integral components of the cell in that they regulate the expression of important genes through the status of their acetylation. However, among them, HDAC6 demonstrates predominant cytoplasmic roles – chaperone activation, tubulin acetylation and clearance of protein aggregates through aggresome formation ^8^. Ding et. al. reported that the AD brain shows higher levels of HDAC6 as compared to the healthy brain ^9^. While some studies present HDAC6 inhibitors as therapeutics for AD ^10,11^, others demonstrate its role in the degradation of Tau and mitigation of the effects of Tau in AD ^12–15^. In this perspective, HDAC6 shows great potential as a therapeutic strategy and more studies need to be done in order to achieve a resolution. ^16^ HDAC6 interacts with multiple proteins in the cell, most prominently tubulin and heat shock proteins while also forming a part of various protein networks and signalling pathways in the cell ^17^. We have previously reported that the treatment of neurons with HDAC6 ZnF UBP demonstrates enhancement in the formation of neurites and migratory structures such as the lamellipodia, filopodia and podosome-like structures, potentially regulating actin dynamics in the neuron ^13^. In the present study, we have explored the possible signalling involved in the effect of HDAC6 on the actin dynamics and established a further understanding of how this helps HDAC6 to facilitate the navigation and migration of the neurons. This work paves the way for the discovery of a more efficient therapeutic possibility for neurodegenerative diseases like AD, by utilizing the potential of Histone Deacetylase 6 domain in tackling the closest aspect of neurodegeneration, i.e., damage to the cytoskeleton.

## 2. MATERIALS AND METHODS

### 5.1 Chemicals/Reagents

Luria Bertani broth [Himedia]; Kanamycin, Imidazol, NaCl, Phenylmethylsulfonylfluoride (PMSF), APS, DMSO, Ethanol (Mol Bio grade), Isopropanol (Mol Bio grade) were purchased from MP biomedicals; IPTG and Dithiothreitol (DTT) from Calbiochem; Terrific broth, MES, BES, SDS and TritonX-100 from Sigma; EGTA, Protease inhibitor cocktail, Tris-base, 40% Acrylamide, TEMED from Invitrogen. For cell culture studies, Advanced Dulbecco Modified Eagle’s Medium (DMEM), Fetal Bovine Serum (FBS), Horse serum, Phosphate Buffered Saline (PBS, cell biology grade Gibco), Trypsin– EDTA, Penicillin–Streptomycin, RIPA buffer, DAPI and Prolong Diamond antifade mounting media were also purchased from Invitrogen. 18 mm coverslip was purchased from Bluestar for immunofluorescence. In immunofluorescence (IF) and western blot (WB) experiments, the following antibodies were used: F-actin (Phalloidin Alexa 488; A12379) [1:40 dilution for IF], TKS5 (Rabbit; PA5-58169) [1:100 for IF, 1:1000 for WB], β-actin (Mouse; MA515739) [1:2000 for WB], Arp2 (Rabbit; PA5-27879) [1:100 for IF, 1:1000 for WB], WASP (Rabbit; PA5-99392) [1:150 for IF], Goat anti-mouse IgG (H+L) secondary antibody HRP (32430) [1:1500], Goat anti-rabbit IgG (H+L) cross-adsorbed secondary antibody HRP (A16110) [1:10000], Goat anti-rabbit IgG (H+L) Cross adsorbed secondary antibody with Alexa Fluor 555 (A-21428) [1:500] all purchased from ThermoFisher Invitrogen.

### 5.2 Preparation of HDAC6

The cloning of HDAC6 ZnF UBP was done in pT7C plasmid and the transformation in BL21* cells. Following primary and secondary inoculation, the culture was induced, at O.D. (600 nm) = 0.6, with 0.2 mM IPTG at 16°C, 180 rpm. Induced cells were harvested after 14 hrs and lysed using lysis buffer in the homogenizer (Constant cell disruption system) at 15 kPSI. The lysate was centrifuged at 40,000 rpm for 40 mins to separate the debris. Purification was performed in two stages: Ni-NTA affinity chromatography followed by size-exclusion chromatography. The supernatant containing the protein was incubated with the Ni-NTA beads for 1.5-2 hours for binding. For the affinity chromatography, 20 mM Imidazole was added to the buffer (Tris-CL pH 8.0) for column wash, while 1000 mM Imidazole for the elution. The high concentration of Imidazole was removed through dialysis with Tris-Cl buffer containing no Imidazole. The dialyzed sample was concentrated to be loaded for size-exclusion (column: Superdex 75 pg Hi-Load 16/600). UV absorbance was monitored at 214 and 280 nm and the fractions corresponding to the absorbance peak were collected. Fractions were pooled, concentrated (5 kDa centricon) and the final protein concentration was estimated through absorbance at 280 nm as well as the Bicinchoninic acid assay.^12,13,18^

### 5.3 Neuronal cell culture, immunofluorescence and colocalization analysis

The cell line used for the study was Neuro2a (N2a; passage 10-20; ATCC CCL-131), maintained in Advanced DMEM supplemented with 10% FBS (Fetal Bovine Serum), 1X Penicillin-Streptomycin, 1X L-glutamine and 1X Anti-Anti (antifungal). For the immunofluorescence staining, 2 x 10^4^ cells were seeded on an 18 mm coverslip in a 12-well culture plate. After the cells were attached to the substratum (24 hrs), the complete media was replaced with the serum-starved (0.5% FBS) media and HDAC6 ZnF UBP (50 nM) was added. The cells were allowed to incubate with HDAC6 for 24 hours after which they were fixed using 4% PFA (paraformaldehyde) for 20 mins at R.T. The fixed cells were washed with 1X PBS and the coverslip was blocked with 5% blocking buffer (5% Horse Serum + 0.2% Triton X-100, detergent in 1X PBS). The coverslip was incubated with the primary Ab (prepared in the blocking buffer) overnight at 4°C. The coverslip was washed to remove any unbound or non-specifically bound Ab. The washed coverslip was further incubated with the secondary Ab for 1 hr at R.T., followed by washing. The coverslips were incubated with DAPI nuclear counterstain for 5 mins, followed by inverted mounting on glass slides for visualization under the epifluorescence microscope (Zeiss Axio observer 7.0 Apochrome 2.0) and confocal microscope (Leica microsystems LASX 4.4.0.24861). The images were captured at 63X magnification (oil immersion) in epifluorescence and at 100X in confocal. The intensities of each protein and their cellular localization were studied through the quantification analysis of the images in the Zen 3.0 and the LASX 3.5 softwares. For a 3D analysis of the colocalization, orthogonal images were acquired for each group, where a series of images was captured at every plane/slice. The main article has representative images from the confocal microscopy, while the epifluorescence images can be found in supplementary figure S1A and the corresponding orthogonal in fig. S1B.

### 2.4 Western blotting for protein expression

30 x 10^4^ N2a cells were seeded on 6-well culture plates and after cell attachment, media was replaced with serum-starved media and HDAC6 ZnF UBP was added for the treatment. After 24 hours of treatment, the cells were harvested (0.05% Trypsin-EDTA) in 1X PBS (to avoid any interference from the serum proteins in the media). The cell pellets were lysed using RIPA buffer (with 0.2% Triton X-100) and the total protein concentration was determined using Bradford’s protein estimation. Accordingly, 75 µg of protein (cell lysate) was loaded on the SDS-PAGE gel (10% resolving for Arp2 and WASP; 6% resolving for TKS5) and electrophoresis was carried out at 90V. The blot membrane was then blocked using 10% blocking buffer (milk powder in 1X PBST; 1X PBST = 1X PBS + 0.1% Tween-20), to avoid any non-specific binding of the antibodies. The blot was incubated with the primary antibody overnight at 4°C, followed by the secondary antibody (Horse Raddish Peroxidase tagged) for an hour at R.T. The blot was washed with 1X PBST after each antibody incubation, for the removal of unbound antibodies. The protein bands are visualized using the developing solution (ECL substrate + Clarity solution, 1:1), applied on the blot and developed in the developing unit. The protein bands are quantified based on their intensities. The ratio of the intensities of the target protein and the loading control bands determines the level of that protein in the cells (control *vs.* treated). The whole blots for each protein can be found in supplementary figure S2: A) TKS5, B) Arp2 C) WASP D) β-actin.

### 5.5 Wound healing assay

50 x 10^4^ cells (N2a neuron and N9 microglia CVCL-0452) were seeded on 6-well culture plates. After the cells had settled, wounds were created in the wells using a tip, followed by a PBS wash to clear the disturbed cells from the wound. HDAC6 ZnF UBP was added in the serum-starved media to the wells. 50 µM ATP was used as the inducer for migration. 0 and 24 hrs images of the wounds were captured in a bright field microscope (10X magnification). The wound diameter is noted at three different points across the wound lines. ^7,19,20^

Wound closure/healing was calculated as:

% Wound closure = (Wound diameter at 24 hrs/ Wound diameter at 0 hr) x 100

### 5.6 Trans well migration assay

The assay was performed in a 24-well transwell plate with inserts (0.5 µm pore membrane). The treatment media was added to the base of the well, while the cell suspension (0.5 x 10^4^) was added to the insert. After 24 hours, the inserts were subjected to fixation with 4% PFA. The inserts were then placed in a primary stain (0.2% crystal violet) followed by washing of the unbound stain with PBS. The upper/inner side of the insert is swabbed leaving only the cells that have migrated to the other side of the membrane. Bright-field images were captured for the stained cells on the inserts (10X and 20X magnification) and counted using the particle counter plugin of FIJI (Fiji is just ImageJ) ^7^.

### 5.7 Statistical analysis

For all the studies, assays were performed in triplicate sets. The statistical validation for all the quantitative data was performed using unpaired t-test and plotted using Tukey’s analysis at 99% confidence interval. For the immunofluorescence intensity and colocalization quantification, more than 25 cells (across different fields) were analyzed. For the wound healing assay, around 25 fields were considered. For the trans-well migration assay, around 20 fields were considered. The plotting and significance analysis were done using GraphPad Prism 8.0 application and the p-values were assessed as follows: p-value≥0.01 implies n.s. (non-significance); the data is considered significant when p<0.01 (p<0.01 results in *, p<0.001 in ** and p<0.0001 in ***).

## 3. RESULTS

HDAC6 has been widely reported to have interactions with numerous cytoskeletal proteins. We therefore asked whether HDAC6 would be involved in one of the important cytoskeletal phenomena – migration. Fig. 1A represents the hypotheses of our present study – i) the role of HDAC6 in migration, ii) whether this migration is facilitated by actin-rich structures called podosomes, and consequently, iii) the role of actin remodelling in neuronal migration. We were interested in understanding the effect of extracellular HDAC6 ZnF UBP treatment to Neuro2a cells, on the actin branching activity through studying changes in Arp2 localization and expression, actin polymerization through WASP and podosome formation through monitoring TKS5 changes.

**Figure 1.**
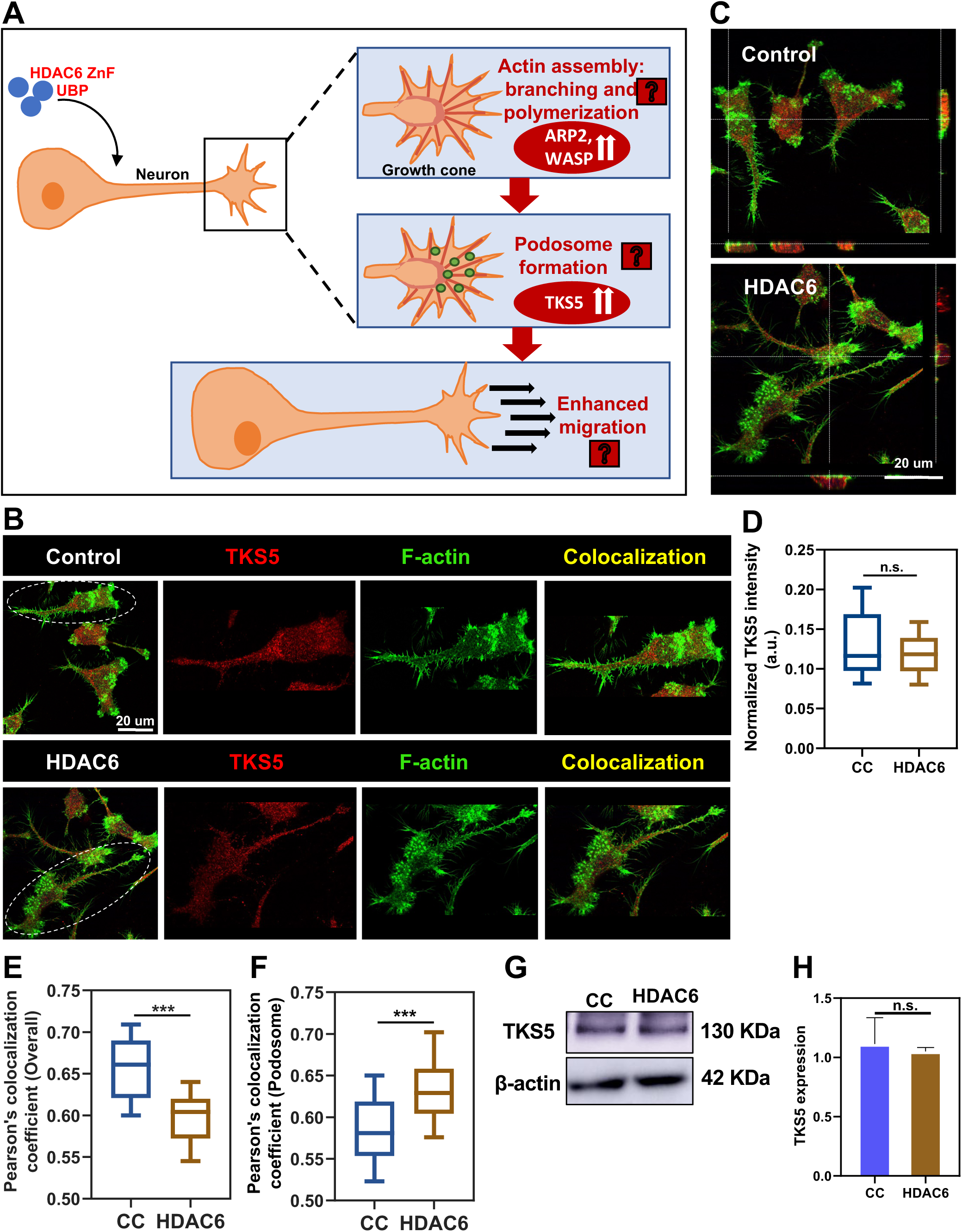
Actin-rich punctae on neuronal cells are podosomes. A) Schematic denoting the hypothesis of the study – HDAC6 ZnF UBP treatment to neurons results in: increased actin branching (Arp2 protein), increased podosome formation (TKS5 protein) and increased migration. B) Confocal images with emphasis on a single cell from the field, showing localization of F-actin (green), TKS5 (red) and their interaction (colocalization) in the merge (DAPI – blue, Merge - yellow). Podosome-specific region and and one with no podosomes have been magnified and highlighted (white dotted circles). C) The orthogonal projections depict the foci of actin and TKS5 colocalization in the control (CC) and HDAC6 ZnF UBP treated (HDAC6) cells. D) Fluorescence intensity quantification shows no significant difference in TKS5 levels between the CC and HDAC6-treated cell groups (Significance n.s. p>0.01 n=3 sets) E) Plot showing the Pearson’s coefficient of colocalization between TKS5 and F-actin in the entire cells (podosome and non-podosome regions included termed as overall colocalization), depicts higher colocalization in the cell control group. (Significance *** p<0.0001; n=3 sets, 62 cells) F) Plot showing the Pearson’s coefficient of colocalization between TKS5 and F-actin only in the podosome structures, depicts a higher level of colocalization in the treated groups. (Significance *** p<0.0001; n=3 sets, 39 cells) G) Western blot analysis of TKS5 with β-actin as loading control also agrees with the immunofluorescence protein expression analysis and shows no significant change among the cell groups. H) Quantification of the western blot analysis showing that the difference in TKS5 expression between the control and treated groups is not significant (p>0.01; n=3 sets).

### 2.1 Podosome formation is enhanced in the presence of the HDAC6 ZnF UBP domain

Podosomes are actin-rich structures characteristic of cells of the migratory nature. ^6^ Formation of podosome structures involves the recruitment of intermediate molecules (like cortactin) required for actin assembly and remodelling. ^16^ TKS5 is an important adaptor molecule in this recruitment process and is abundantly present in the podosomes making it a largely used marker for podosomes in cells. ^21–23^ Neuro2a cells were extracellularly treated with HDAC6 ZnF UBP and mapped for F-actin and TKS5 localization. The distinct actin-rich punctae observed in both groups showed the presence of TKS5 and were thus attributed as podosomes. Figure 1B shows the localization and colocalization of TKS5 and F-actin in the control and HDAC6 treated cells Fig. 1C highlights the colocalization points in the cells, through an orthogonal representation of the cells. On quantification of the colocalization in the whole cell versus the podosome regions, it was found that the HDAC6-treated cells showed higher F-actin-TKS5 colocalization in the podosomes (only podosome area) (Fig. 1F), whereas the control groups showed higher whole cell colocalization (whole cell including podosome and non-podosome areas) (Fig. 1E). The protein expression levels remain unchanged on treatment, as seen in the fluorescence intensity quantification plot (Fig. 1D), and confirmed through the western blot analysis (Fig. 1G, H). Thus, it can be inferred that the distinct actin-rich structures we have reported in these neuronal cells are indeed podosomes. Moreover, the presence of HDAC6 ZnF UBP modifies the localization of TKS5 to the podosomes, thereby helping in their regulation.

Podosomes are dynamic structures, found in different stages or classes in cells (Fig. 2A). They start by forming distinct single podosomes, moving on to forming clusters among themselves. These clusters attain specific structures such as a ring structure, which are termed rosettes, and the last stage is when the podosome clusters decorate the periphery of the cells and develop into podosome belts. ^4,24^ To understand this phenomenon, we performed a time-dependent immunofluorescence analysis for the colocalization. The cells were treated with HDAC6 ZnF UBP for 1, 3, 6, 12 and 24 hrs and mapped for TKS5 and F-actin. Fig. 2B shows merged images for the colocalization of TKS5 and F-actin at each time point, with emphasis on the whole field (inset) as well as single cells, with the corresponding intensity plot in Fig. 2C. A significant decrease was seen in the intensity of TKS5 at 6 and 12 hrs of HDAC6 treatment, although the mechanism has not been explored. Figure 2D shows a comparison of the number of cells bearing podosomes across the different time points and does not show significant changes with time. There was no significant difference in the classes of podosomes across different time points (Fig. 2G-J), however, there was an increase in TKS5-actin colocalization in the podosomes, with increasing time of HDAC6 treatment, as compared to the control (Fig. 2F). It is interesting to note, however, that the whole-cell colocalization showed comparatively contrasting results, i.e., the control group showed higher colocalization than the treated (Fig. 2E). Thus, it can be inferred that the effect of HDAC6 is not on the protein levels or overall podosomes but is in fact on the cellular localization of TKS5 and the point of colocalization of the proteins.

**Figure 2.**
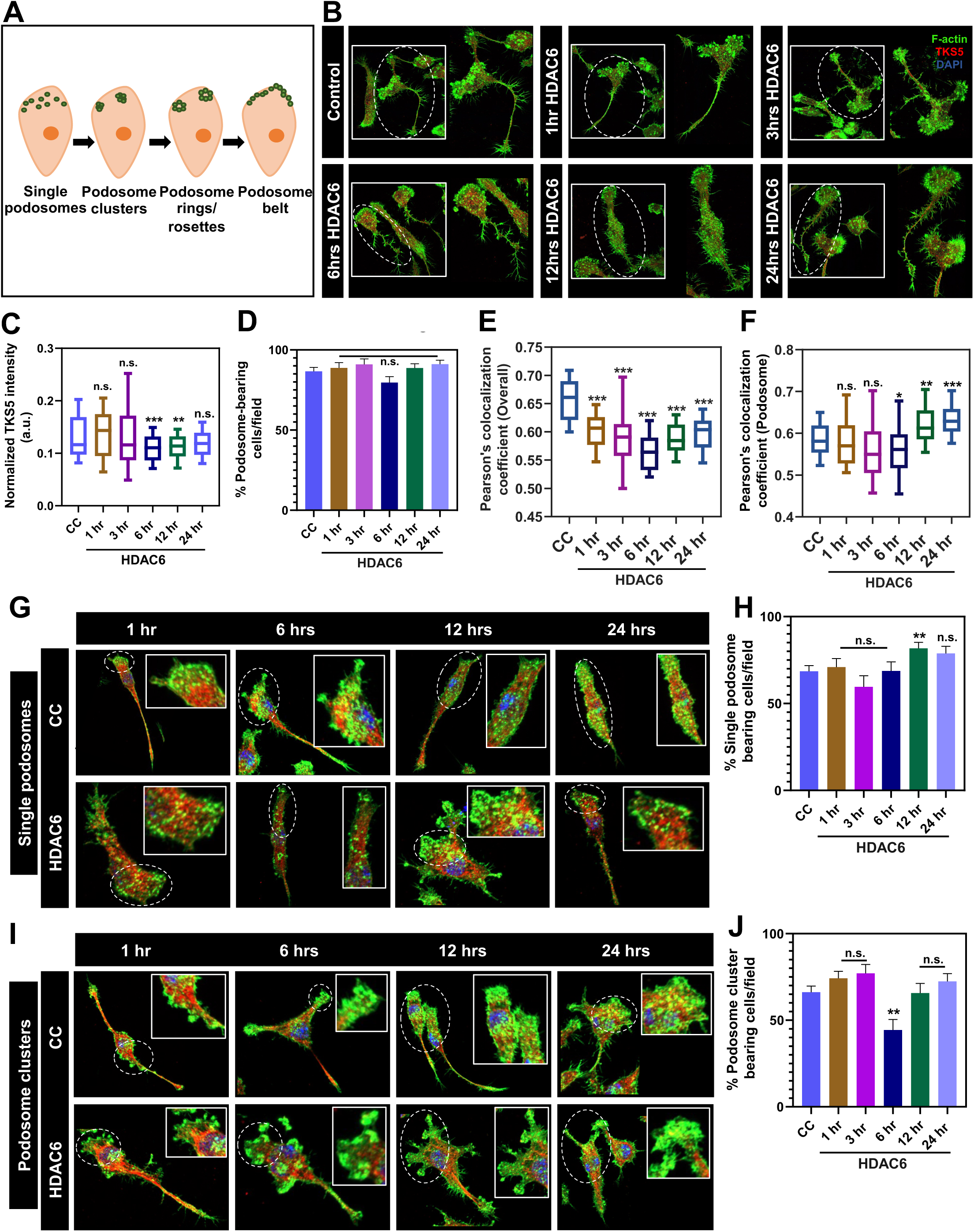
Time-dependent study on podosomes formation with HDAC6. A) Stages of podosome formation. The first stage is that of individual actin-rich punctae termed single podosomes. They come together to form podosome clusters. In the mature stages, podosome clusters arrange in either a ring formation, known as rosettes or at the periphery of the cell, called podosome belts. B) N2a cells were subjected to treatment with HDAC6 ZnF UBP for 1, 3, 6, 12 and 24 hours, followed by fixation and staining with anti-TKS5 antibody (red) and Phalloidin (green; F-actin). The images show an enlarged single cell with an inset of the whole field (the enlarged single cell is marked in the white dotted circle) C) Fluorescence intensity of TKS5 at different time points shows no significant change in the TKS5 expression. (Significance is n.s. p>0.01; n=3 sets) D) The number of cells containing podosomes per field was counted and plotted as the percentage of cells with podosomes (% Podosome-bearing cells), revealing no significant changes across time points; (Significance is n.s. p>0.01; n=3 sets) E) The overall TKS5-Factin colocalization demonstrates a significantly lower value in treated cells compared to the control. Significance was calculated for each time-point of the treated groups with respect to the control (n.s.= non-significant, p>0.01; * = p<0.01; ** = p<0.001; *** = p<0.0001; n=3 sets, 36 cells),the most significant being the 6 hr treatment. F)The plot for ‘podosome’ colocalization demonstrates significantly higher values in the treated cells compared to control. Significance was calculated for each time-point of the treated groups with respect to the control (n.s.= non-significant, p>0.01; * = p<0.01; ** = p<0.001; *** = p<0.0001; n=3 sets, 36 cells). G) Single podosomes were observed in the control as well as the treated groups. H) No. of cells containing only single podosomes were counted and plotted as the % cells with single podosomes, revealing no significant changes across time points (Significance is n.s. p>0.01; n=3 sets). I) Podosome clusters were observed in the control as well as the treated groups. J) The no. of cells containing podosome clusters was counted and plotted as the percentage of cells with podosome clusters, demonstrating no significant change across time, except in 6hrs HDAC6 treatment. (Significance is n.s. p>0.01; n=3 sets).

### 2.2 HDAC6 facilitates increased actin branching in podosomes

To prove our proposition regarding the role of HDAC6 in actin remodelling, the effect of HDAC6 was studied on the branching activity of actin through immunofluorescent mapping of the actin regulatory protein ARP2 ^25–28^. ARP2 is a part of the Arp2/3 complex, that facilitates actin branching or the formation of new actin filaments on the old ones (Fig. 3A). As can be seen in Fig. 3B, Arp2 and F-actin colocalize in the neurons, both in the podosome regions and the extensions.Fig. 3D presents the orthogonal images for the colocalization points between Arp2 and F-actin. Analysis of colocalization was performed similarly to TKS5 and there was a significant difference observed between the colocalization foci of the control and treated cells – HDAC6 treatment confers lower whole cell colocalization (fig. 3E) but higher podosome colocalization (fig. 3F), with the reverse observed in control cells. The protein levels remained unchanged on treatment (Fig. 3C), confirmed by the western blot analysis (Fig. 3G, H). Hence, it can be inferred that in the presence of HDAC6 ZnF UBP, there is increased interaction between Arp2 and F-actin, promoting actin branching at the site of podosomes.

**Figure 3.**
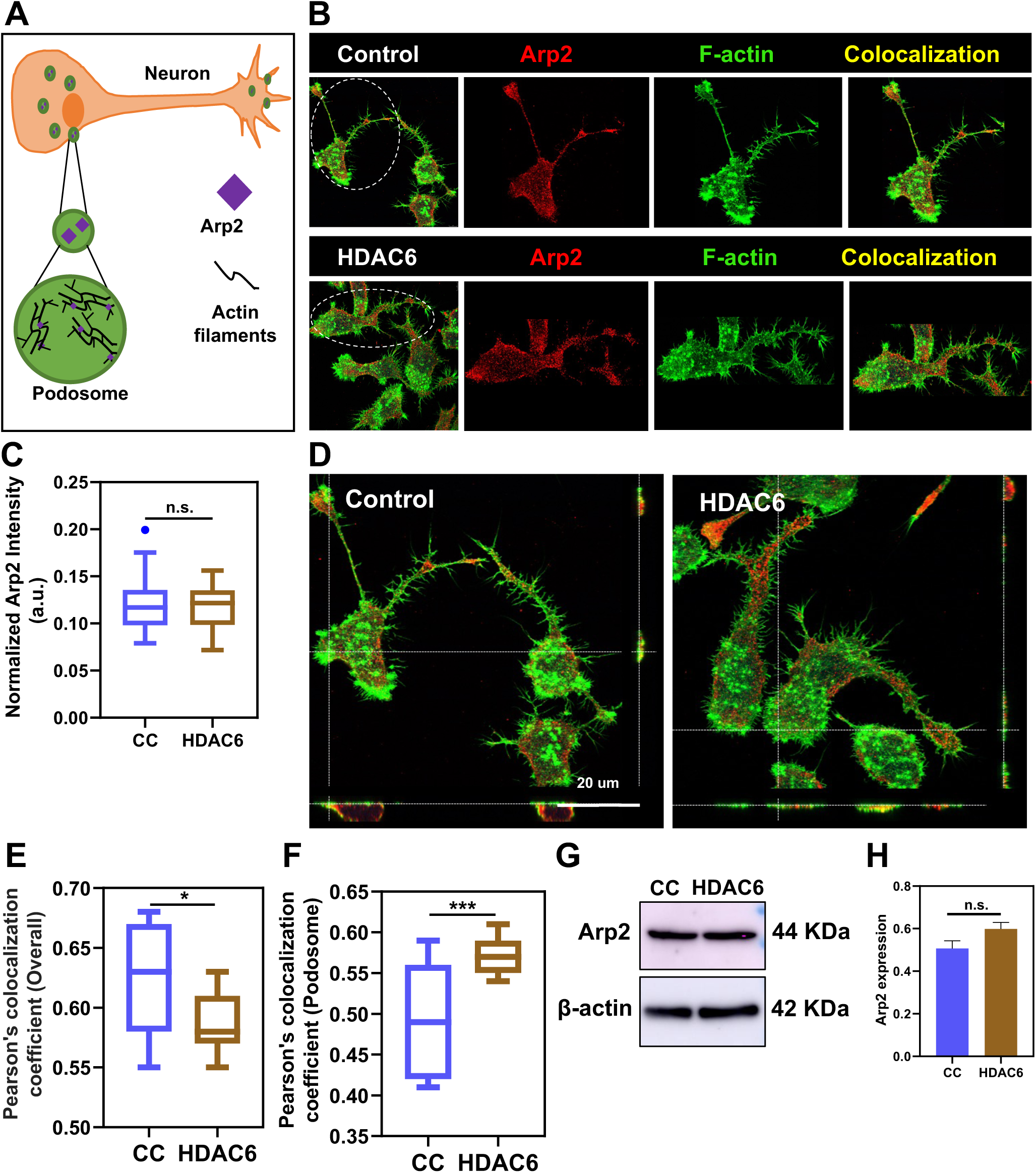
HDAC6 enhances actin branching in podosomes. A) Scheme showing the involvement of Arp2 in podosome formation. In podosomes, Arp2 facilitates the branching of actin filaments, which results in the complex cytoskeletal networks and structure of the podosomes. B) Confocal images with emphasis on a single cell from the field, showing localization of F-actin (green), Arp2 (red) and their interaction (colocalization) in the merge (DAPI – blue, Merge - yellow). Podosome-specific region and and one with no podosomes have been magnified and highlighted (white dotted circles). C) Fluorescence intensity quantification shows no significant difference in Arp2 levels between the groups (Significance n.s. p>0.01 n=3 sets, 36 cells). D) The orthogonal projections depict the foci of actin and Arp2 colocalization in both cell groups. E) Plot showing the Pearson’s coefficient of colocalization between Arp2 and F-actin in whole cells (overall) depicts higher colocalization in cell control group (Significance * p<0.01 n=3 sets, 19 cells). F) Plot showing the Pearson’s coefficient of colocalization between Arp2 and F-actin in the podosome structures in cells, depicts higher colocalization in the HDAC6 treated group (Significance *** p<0.001 n=3 sets, 15 cells) G) Western blot analysis of Arp2 with β-actin as loading control also agrees with the immunofluorescence protein expression analysis and shows no significant change among the cell groups. H) Quantification of the western blot analysis showing that the difference in Arp2 expression between the control and treated groups is not significant; n.s. p>0.01 n=3 sets.

### 2.3 HDAC6 is involved in the actin nucleation process in podosomes

We have previously hypothesized that HDAC6 potentially activates Cdc42, Rac and thus the WASP (Wiskott-Aldrich Syndrome Protein) and WAVE (WASP-Associated Verprolin homolog Protein) complexes ^16^. The WASP and WAVE proteins are very important in that they are responsible for the activation of the ARP2/3 complex. We studied the effect of HDAC6 on the WASP localization within the podosomes. WASP was seen to colocalize with F-actin similar to Arp2 (Fig. 4A, C). The quantification analysis (Fig. 4D, E) showed that HDAC6 confers higher colocalization between WASP and F-actin in the podosomes rather than the whole cell, compared to control cells. The protein expression levels showed a significant difference in the immunofluorescence analysis (Fig. 4B), however western blot analysis revealed a non-significant difference in the WASP expression levels (Fig. 4F, G). It can therefore be suggested that HDAC6 participates in the processes leading to the remodelling of actin for the formation of podosomes.

**Figure 4.**
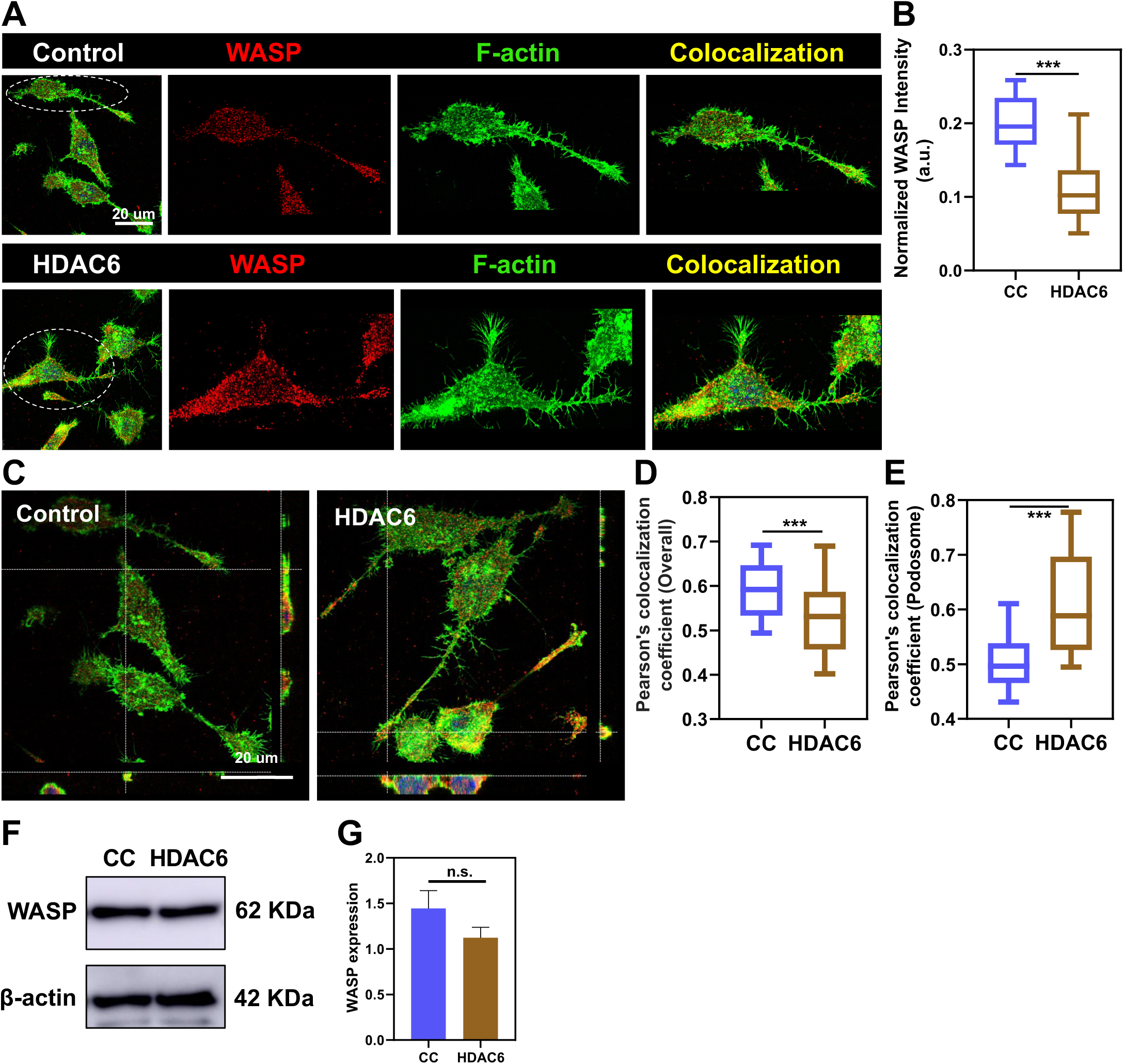
HDAC6 facilitates WASP-mediated actin nucleation in podosomes. A) Confocal images with emphasis on a single cell from the field, showing localization of F-actin (green), WASP (red) and their interaction (colocalization) in the merge (DAPI – blue, Merge - yellow). Podosome-specific region and and one with no podosomes have been magnified and highlighted (white dotted circles). B) Fluorescence intensity quantification shows a significant difference in the levels of WASP between the control and treated groups (Significance *** p<0.001 n=3 sets). C) The orthogonal projections depict the foci (magnified in podosomes and cell regions) of actin and WASP colocalization in the control and HDAC6 ZnF UBP treated cells. D) Plot showing the Pearson’s coefficient of colocalization between WASP and F-actin in whole cells (overall), depicts higher colocalization in the cell control group (Significance *** p<0.001 n=3 sets, 43 cells). E) Plot showing the Pearson’s coefficient of colocalization between WASP and F-actin in the podosome structures in cells, depicts higher colocalization in the HDAC6 treated group (Significance *** p<0.001 n=3 sets, 20 cells). F) Western blot analysis of WASP with β-actin as loading control with the two groups. G) Quantification of the western blot analysis shows a non-significant decrease in WASP expression in HDAC6 treated cells, compared to the control group (Significance n.s. p>0.01 n=3 sets).

### 2.4 HDAC6 potentially enhances podosome-mediated migration in neurons

From the above findings, it could be deduced that HDAC6 has a role to play in the actin remodelling machinery of the cells. Thus, we hypothesized that this domain would consequently take part in the cytoskeletal phenomena of migration since it is the major biological function of podosomes. We noticed that the extracellular treatment with HDAC6 ZnF UBP domain resulted in an activated migratory morphology in cells, *i.e.,* increased formation of the cytoskeletal structures of filopodia and podosomes in the cells (Fig. 5A). The number of cells exhibiting the migratory morphology in both the cell groups were determined and it was found that the HDAC6 treated cells showed significantly less normal morphology and more migratory morphology (Fig. 5B). Fig. 5C emphasizes the cytoskeletal structures of podosomes and filopodia, in the gray-scale images of F-actin in cells. The number of filopodia (Fig. 5D) and podosomes (Fig. 5E) were determined per cell and it was found that the HDAC6-treated cells possessed a higher number of migratory structures.

**Figure 5.**
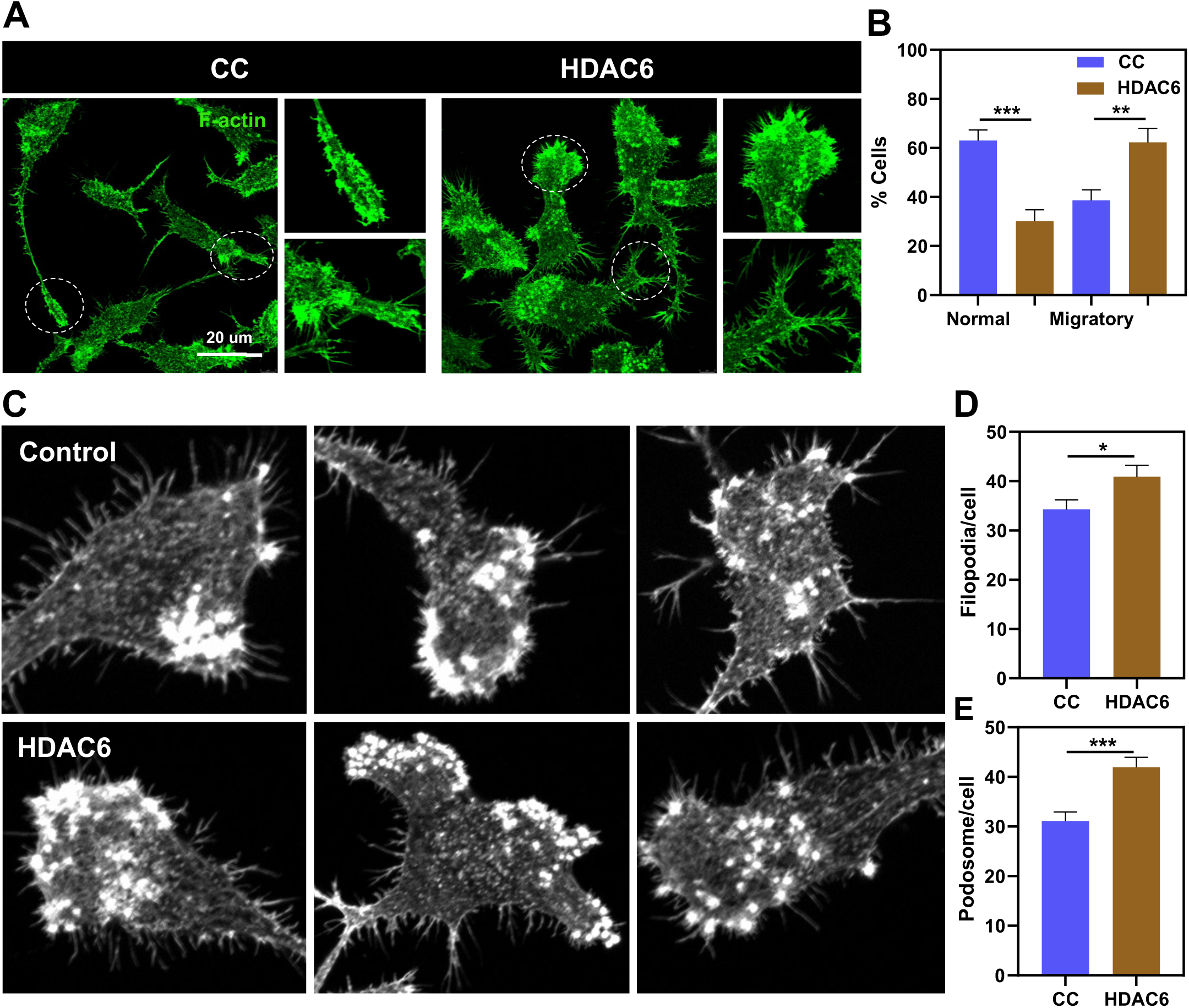
Effect of HDAC6 on neuronal morphology. A) Treated cells displayed a transition to the migratory morphology (encircled in white and magnified). B) No. of cells showing normal and migratory morphology in control (CC) and treated (HDAC6) groups have been ploted as % cells The comparison reveals a higher percentage of normal morphology cells (Significance ** p<0.001, n=8 sets) in control than the HDAC6 group, which shows higher percentage of cells with the migratory morphology (Significance ** p<0.001, n= 8 sets). C) Grayscale images for F-actin structures of filopodia and podosomes in the control and HDAC6 treated cell groups. D) Plot with the number of filopodia per cell shows that HDAC6 treatment confers higher filopodia formation compared to control cells (Significance * p<0.01, n= 3 sets, 64 cells). E) Plot with the number of podosomes per cell shows that HDAC6 treatment confers higher podosome formation compared to control cells (Significance *** p<0.0001, n= 3 sets, 51 cells).

It was therefore interesting to look at the migration capacity of the cells in the presence of extracellular HDAC6 ZnF UBP. This was done using both the wound-healing (2D) and transwell (3D) migration assays. For all the migration assays, microglia were used as the positive control for migratory cells and ATP as the positive control for the induction of migration. In the 2-D wound healing assay, the overall wound closure was observed to be ≈20%, however no significant differences were observed between cell control and HDAC6 treated cells (Fig. 6A, B). On the other hand, the microglia demonstrated comparatively higher wound closure (∼25%), but similarly, no significant difference was observed between the cell control and HDAC6-treated cells (Fig. 6C, D). In the 3-D trans-well migration assay, an increase in neuronal migration was observed in the case of HDAC6 treatment (∼70 cells/field), compared to the control group (∼50 cells/field). Like the wound healing assay, ATP showed no effect on the migration of neurons (Fig. 6E, F). Microglia followed a similar pattern to that of its wound healing assay - no significant difference between control (∼300 cells/field),HDAC6 (∼300 cells/field) and ATP (∼300 cells/field) (Fig. 6G, H). To summarize, neurons show significant alteration of their 3D migration profile on HDAC6 treatment, whereas, the 2D migration profile remains unaltered.

**Figure 6.**
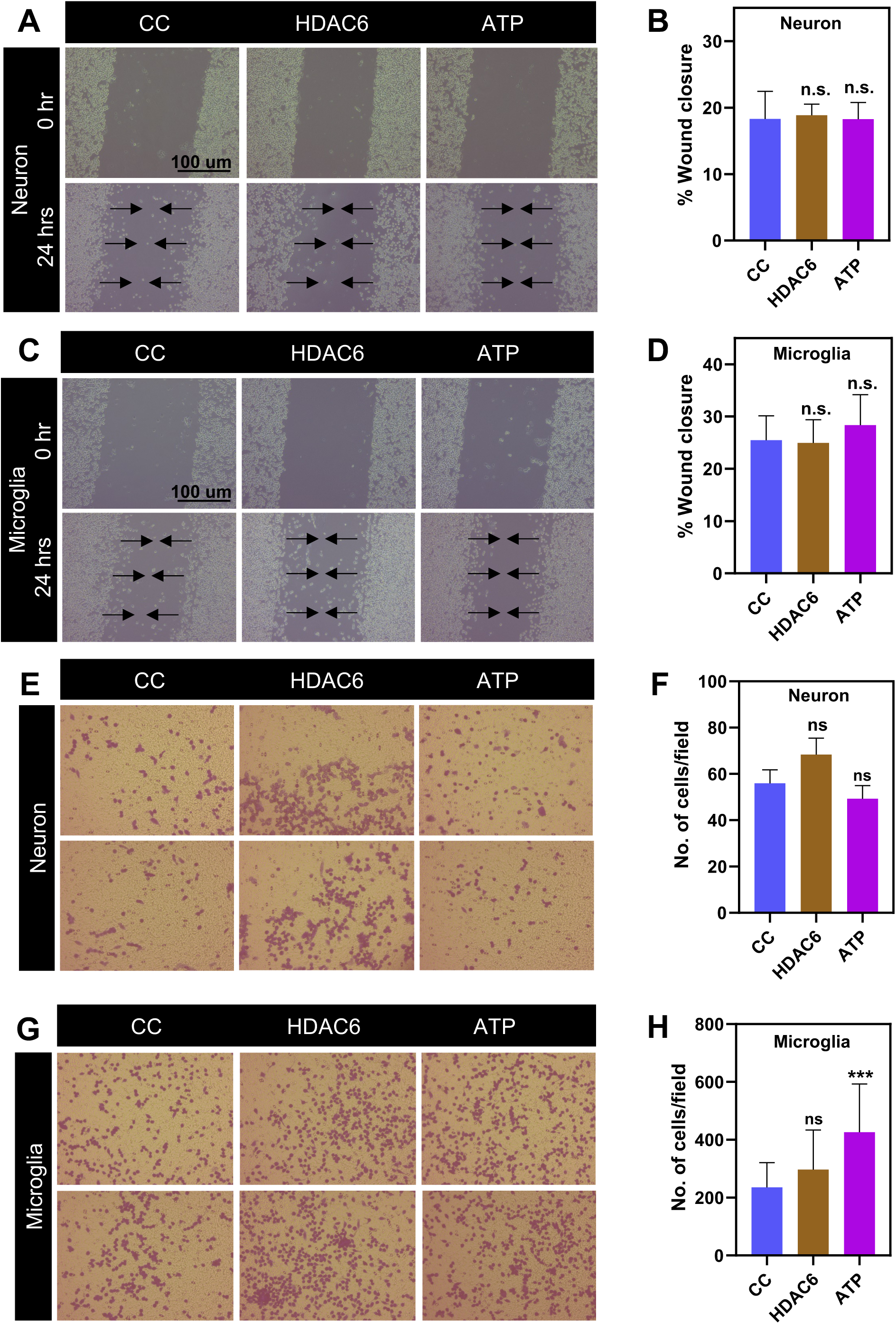
Effect of HDAC6 on neuronal migration. The migration capacity of the cells was tested through wound-healing (2D assay) and transwell (3D) migration assays. Microglia was used as the positive control for migratory cells and ATP the control for the induction of migration. Wound healing assay images are phase contrast images captured at 10X magnification, while transwell migration images are from brightfield microscopy at 20X. A) For the wound healing assay, neuronal cells (Neuro2a) were allowed to form a dense cell monolayer and wounds were created at 3 positions, followed by treatment with HDAC6 ZnF UBP. Images of the wound, captured at 0 and 24hrs of treatment reveal the difference in the wound diameter between the 2 time points; the arrows represent the direction of migration of cells towards closure of the wound. B) The percentage of wound closure was plotted for different fields, based on the difference in wound diameters at the initial and final treatment time points. Plot suggests no significant difference in the migration of the 3 groups of cells. (Significance is n.s. p>0.01; n=3 sets). C) The wound healing assay for microglial N9 cells; images captured at 0 and 24 hrs. D) Percent wound closure in microglia shows that HDAC6 shows slightly lower wound healing capacity, while ATP, as the positive control, shows an increase in migration, compared to the control group. (Significance is n.s. n=3 sets) E) The trans-well migration assay for neuronal cells shows more migration in the case of HDAC6 treatment. F) The number of cells that migrated to the other side of the membrane per field was plotted. In neuronal cells, HDAC6 treatment showed a significantly higher number of migrated cells, compared to the control and ATP groups. (Significance is * p<0.01; n=3 sets) G) The trans-well migration in microglia shows slightly more migration in the treated groups of cells. H) Plot suggests no significant increase in both the HDAC6 and the ATP groups (ATP, HDAC6: Significance n.s. p>0.01; n=3 sets).

## 4. DISCUSSION

The cytoskeleton is an integral part of the function of neurons. Neuronal extensions, neurites, growth cones ^2,29^, podosomes ^5^ are cytoskeletal structures involved in varied neuronal activities - from forming a communication network within the body through synapses, facilitating migration and navigation of the neurons ^30^ to maintaining the overall neuronal homeostasis ^31^. The current study focusses on the effect of neuronal exposure to the ZnF UBP domain of HDAC6 on the cytoskeletal regulation in neurons. This has been done through studies on the actin remodelling machinery in complex cytoskeletal structures of the podosomes. Podosomes have been described as hotspots of adhesion, migration and invasion ^4^ in several cells of the migratory nature. They have been observed and studied in osteoclasts ^32–35^, transformed cells, immune cells ^36^ as well as microglia ^5,7,37^. They are rich in matrix metalloproteinases, which facilitate degradation of the extracellular matrix, thereby enabling migration and navigation of the cells in the extracellular space. Podosome formation is a highly dynamic process involving the major cytoskeletal proteins – actin, tubulin ^36^ and their associated proteins such as talin, Arp complexes, kinases, etc. ^5,37,38^). The confirmation of the presence of podosomes in neurons was performed through the detection of an abundant protein found abundantly in the podosome, TKS5 ^39,40^. TKS5 is a substrate for the receptor tyrosine kinase (Src family of tyrosine kinases), with domains enabling interaction with the plasma membrane. It acts as an adaptor protein, recruiting proteins such as N-WASP and cortactin, required for actin remodelling and podosome formation and maturation ^21–23^. We have observed colocalization of TKS5 with F-actin in the podosome punctae in both the control and treated groups of cells, in agreement with previous reports for different cell types like neural crest cells ^41^, osteoclasts ^42^, fibroblasts, endothelial and adenocarcinoma epithelial cells ^43^. Here, through a differential colocalization analysis (whole cell *vs.* podosome region), we report that HDAC6 ZnF UBP facilitates increase of TKS5 localization in the podosome structures, without any significant change in the expression levels of the protein. This finding is reinforced in the time dependent studies, where we observed increased podosome localization of TKS5 with increasing period of HDAC6 treatment.

The major event in podosome formation is the branching and polymerization of F-actin filaments, carried out by the Arp2/3 complex of proteins ^44^. Arp2/3 proteins are activated either by the Nucleation Promoting Factors such as WAS protein or other actin-binding proteins like cortactin ^16,45,46^. We have shown here that HDAC6 is able to enhance the localization of Arp2 protein in the podosomes, suggesting the involvement of HDAC6 in enhancing branching and polymerization of actin for the regulation of podosome formation ^28^. It is important to note that since Arp2 is involved in actin nucleation, its colocalization with F-actin is precedented; however, the significance lies in the degree of colocalization in the podosome structures between the two cell groups. Further, colocalization analysis of WASP and F-actin also showed higher interaction in the podosomes, in the presence of HDAC6 ZnF UBP. This led us to establish that HDAC6 is possibly involved in WASP-mediated activation of Arp2.

The effect of HDAC6 ZnF UBP treatment on the morphology of the neurons was one of the most interesting findings of this study. The neuronal cells adopted a more migratory morphology, with ruffling of the membrane, increased extensions and filopodia, etc.^47^. To test this, the migration capacity of the cells was determined as an effect of the HDAC6 domain. Interestingly, the 2D migration, represented by the wound healing assay, showed no significant results whereas the 3D or Transwell migration assay showed a significant increase in the migration of neurons with HDAC6 treatment. This indicates the influential effect of HDAC6 on the invasiveness of neuronal cells over its 2D plane migration. Also, the enhanced ability of neurons for 3D migration in the presence of HDAC6 certainly correlates with the observed podosome formation during its exposure. However, it is also important to note here that the effect of HDAC6 exhibited specificity towards neurons and not microglia, the reason for which is still a subject of experiments. Although further optimization and experimentation are required to reach any significant conclusion, these results provide a potential basis for exploration of the role of HDAC6 in neuronal migration.

## 5. CONCLUSION

To summarize our findings, HDAC6 ZnF UBP treatment to neurons results in better actin branching and polymerization, which enhances the formation of podosomes and other migratory structures (filopodia), thereby changing the morphology of the cells and ultimately conferring neuronal cells with an enhanced ability to migrate (Fig. 7). Our observations, along with other reports ^48,49^, form a foundation for the field of neuronal podosomes as key players in neuronal health and disease.

**Figure 7.**
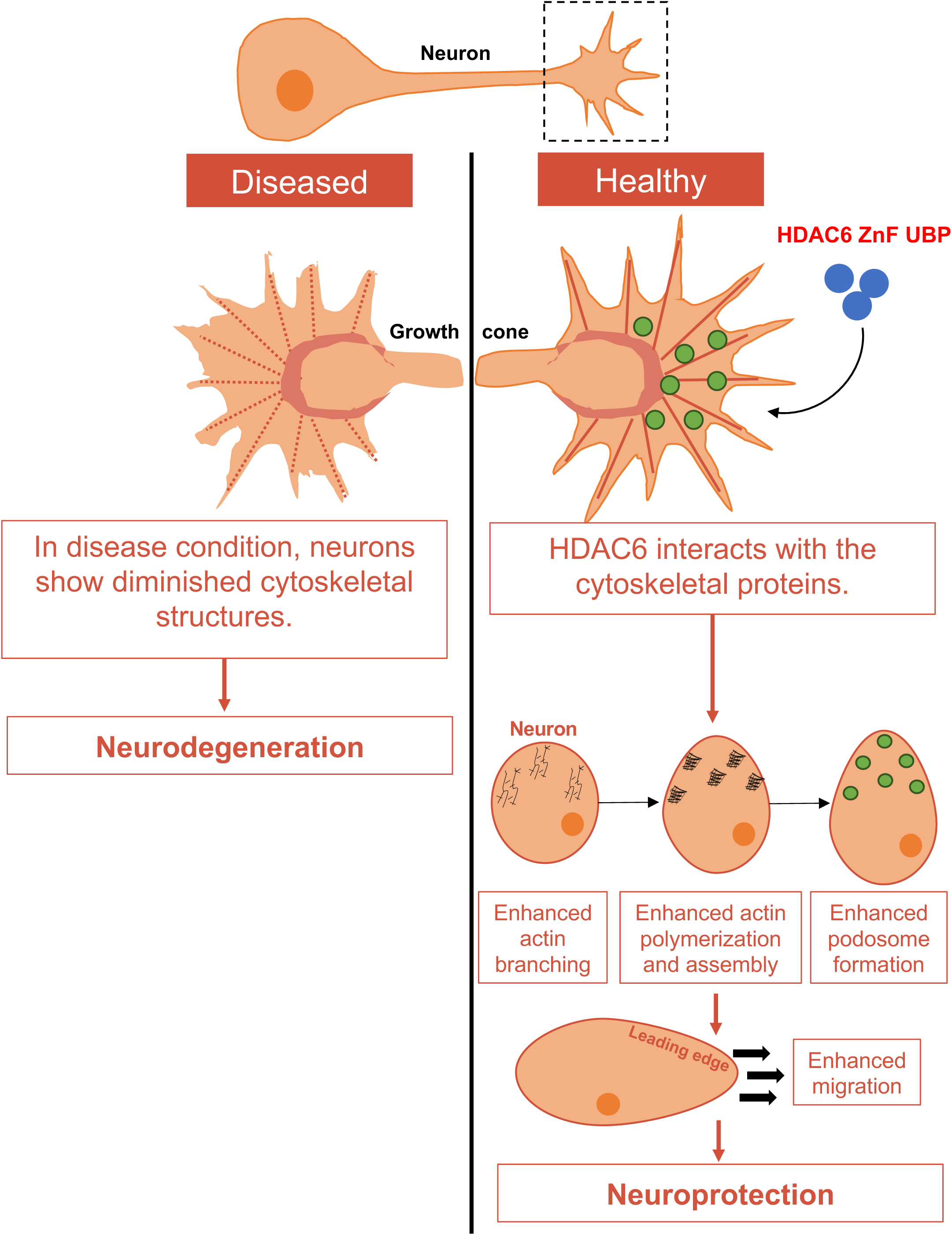
HDAC6 ZnF UBP interacts with the cytoskeleton and confers enhanced migration potential to neurons: A schematic to summarize the findings of the study. The scheme shows a neuron and for cytoskeletal representation, the status of the neuronal growth cone is focused on, in healthy and diseased conditions. In the diseased condition, the cytoskeleton is dysregulated, leading to poor neuronal health and neurodegeneration. On the other hand, on treating cells with HDAC6 ZnF UBP, we propose that HDAC6 interacts with multiple proteins involved in the actin remodelling phenomenon in the neuron. The presence of HDAC6 results in improved actin branching, polymerization and assembly in the podosomes. This contributes towards a change in the morphology of cells giving them a higher ability to migrate and navigate, ultimately conferring protection to the neurons.

### AUTHOR CONTRIBUTIONS

**Tazeen Qureshi:** Investigation, Visualization, Data curation, Formal analysis, Software, Writing-Original draft preparation; **Smita Eknath Desale:** Investigation, Visualization, Data curation, Formal analysis, Software, Writing-Original draft preparation; **Shweta Kishor**

**Sonawane:** Data curation, Formal analysis, Software; **Abhishek Ankur Balmik:** Data curation, Formal analysis, Software; **Hariharakrishnan Chidambaram:** Data curation, Formal analysis, Software, Writing; **Subashchandrabose Chinnathambi:** Conceptualization, Investigation, Visualization, Funding acquisition, Supervision, Project administration, Data curation, Formal analysis, Software, Writing-Original draft preparation, Reviewing and Editing

## ACKNOWLEDGMENTS

We are grateful to Chinnathambi’s lab members for their scientific discussions, helpful suggestions and critical reading of the manuscript. Tazeen Qureshi acknowledges a fellowship from the Department of Science and Technology-Innovation in Science Pursuit for Inspired Research (DST-INSPIRE), India. We also greatly acknowledge internal support from the Department of Neurochemistry, National Institute of Mental Health and Neuro Sciences (NIMHANS), Institute of National Importance, Bangalore.

## DATA AVAILABILITY STATEMENT

The data that support the findings of this study are available from the corresponding author upon reasonable request.

