## Supplementary material for "Extracellular HDAC6 ZnF UBP domain enhances podosome-mediated neuronal migration": SI

**Figure S1**

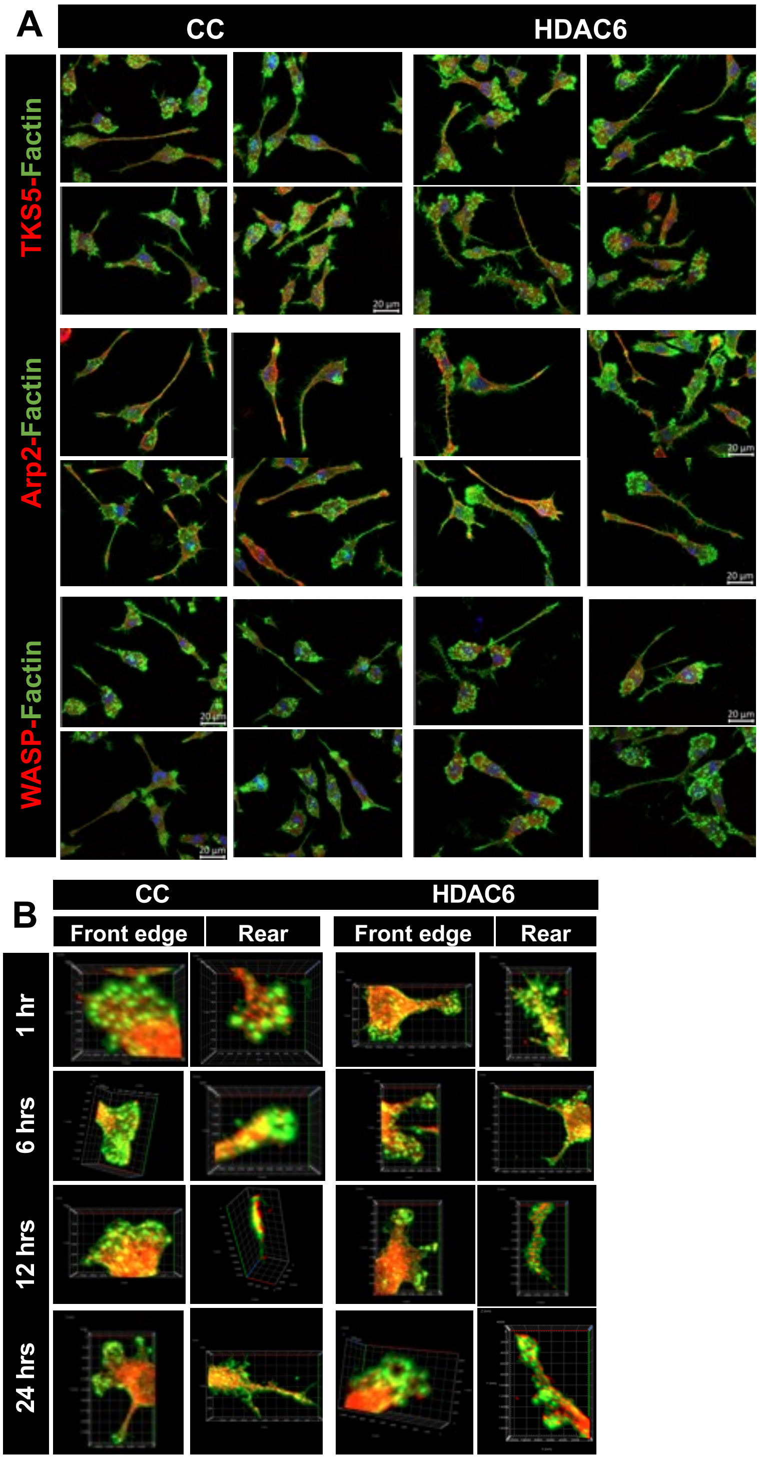


**Fig. S1: HDAC6 ZnF UBP interacts with the cytoskeleton and confers enhanced migration potential to neurons (Epifluorescence microscopy images).** A) A compilation of the representative epifluorescence images for all groups. All the respective quantifications for the particular groups are derived from these images. B) A 3-dimensional view of podosome structures, zoomed in and with emphasis on the colocalization within the podosomes. The front and the rear ends of the cells have been shown to give an overall picture.

**Figure S2**

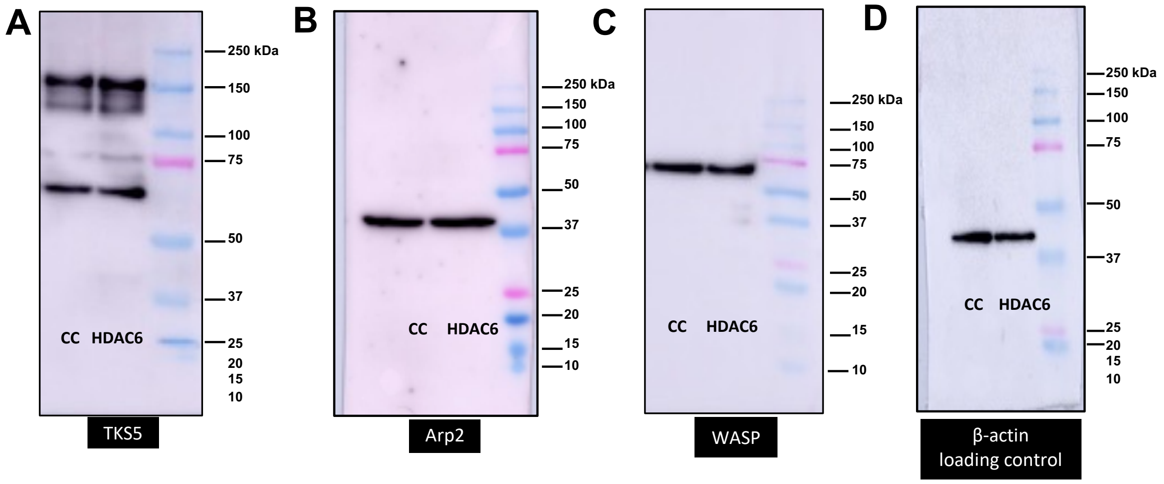


**Figure S2: Expression of cytoskeletal proteins on HDAC6 treatment to cells – Full Western blots.** A) TKS5 at 130 kDa. B) Arp2 at 44 kDa. C) WASP at 62 kDa. D) β-actin at 42 kDa.
